# Estimating dynamic individual coactivation patterns based on densely sampled resting-state fMRI data and utilizing it for better subject identification

**DOI:** 10.1101/2022.01.06.475181

**Authors:** Hang Yang, Xing Yao, Hong Zhang, Chun Meng, Bharat Biswal

**Author notes:** **Correspondence:** Dr. Hang Yang, Chinese Institute for Brain Research, Beijing 102206, China;, Dr. Bharat Biswal, 607 Fenster Hall, University Height, Newark, NJ, 07102, USA.

## Abstract

As a complex dynamic system, the brain exhibits spatially organized recurring patterns of activity over time. Coactivation patterns (CAPs), which analyzes data from each single frame, has been utilized to detect transient brain activity states recently. However, previous CAP analyses have been conducted at the group-level, which might neglect meaningful individual differences. Here, we estimate individual CAP states at both subject- and scan-level based on a densely-sampled dataset: Midnight Scan Club. We used differential identifiability, which measures the gap between intra- and intersubject similarity, to evaluate individual differences. We found individual CAPs at the subject-level achieved the best discrimination ability by maintaining high intra-subject similarity and enlarging inter-subject differences, and brain regions of association networks mainly contributed to the identifiability. On the other hand, scan-level CAP states were unstable across scans for the same participant. Expectedly, we found subject-specific CAPs became more reliable and discriminative with more data (i.e., longer duration). As the acquisition time of each participant is limited in practice, our results recommend a data collection strategy that collects more scans with appropriate duration (e.g., 12~15 mins/scan) to obtain more reliable subject-specific CAPs, when total acquisition time is fixed (e.g., 150 mins). Overall, this work has constructed reliable subject-specific CAP states with meaningful individual differences and provides a starting point for the subsequent applications of individual brain dynamics.

## 1. INTRODUCTION

The brain is a real-time system that relies on dynamic inter-regional coordination to support flexible behavioral and cognitive demands. Accumulated evidence has shown that between-region synchronous oscillations are not static but vary across time (Hutchison et al., 2013; Preti et al., 2017). The time-varying inter-regional interactions can be assessed by dynamic functional connectivity (FC) across sliding windows, and recurring FC patterns have been identified (Allen et al., 2014). An emerging volume-based method, coactivation patterns (CAPs), has also been developed to detect transient brain network dynamics (Liu et al., 2013; Liu & Duyn, 2013). The spatiotemporal dynamics of brain states are supposed to be related to various cognitive processes (Huang et al., 2020; Kupis et al., 2021), demographic characteristics (Murray et al., 2021; Peng et al., 2023) and disorder alterations (Kaiser et al., 2019; Piguet et al., 2021; Rey et al., 2021). The dynamic FC method could only capture slowly varied brain dynamics due to its low sampling rate (i.e., a window lasts for at least 20 s). The CAP method is free from the choice of arbitrary parameters such as window length, and it is capable of detecting transient brain state transitions that happen rapidly. However, traditional CAP analyses are performed at the group level, it is unclear whether reliable subject-specific CAP states can be obtained based on each individual’s own data.

Individual brains are not only heterogeneous in patients (Marquand et al., 2016; Price et al., 2017), but also differ among healthy participants (Mueller et al., 2013). The individual brain differences in structure and function are biologically meaningful (Cui et al., 2020), and are able to predict individual’s behavior (Kong et al., 2019; Sui et al., 2020; Wang et al., 2018) and clinical intervention (Cash et al., 2021; Lynch et al., 2019). The reliability of individual brain features has been validated over time (Gratton et al., 2018; Horien et al., 2019), between datasets (Gordon, Laumann, Adeyemo, et al., 2017) and tasks (Laumann et al., 2015). Additionally, individual brain connectomes are found to be unique to serve as brain fingerprinting for subject identification (Finn et al., 2015; Kumar et al., 2017; Pallares et al., 2018). The identifiability of brain fingerprinting can be manipulated by various cognitive states, increased with development and is hampered in clinical populations (Finn et al., 2017; Kaufmann et al., 2017; Sorrentino et al., 2021). Previous subject identification studies are restricted to connectome constructed by static functional connectivity, while brain dynamics are also meaningful individual features that contribute to identifiability (Menon & Krishnamurthy, 2019). It is unknown whether different CAP states that carrying specific spatiotemporal dynamics are able to distinguish individuals from each other, and whether the identification accuracy can be improved by using individualized CAPs.

In this study, we used a dense sampling dataset, Midnight Scan Club (MSC), to answer the aforementioned questions. Besides the group-level CAPs, we generated the individual CAP states at two levels: leveraging fMRI data from all scans of the subject (subject-level) or based on a single scan of the subject (scan-level). We then assessed the reliability of CAP states, and evaluated the individual differences carried by CAPs using both differential identifiability and identification rate. The contribution of brain regions and networks to subject identification was also estimated. Finally, we explored how individual CAPs varied with different amounts of data, and recommended a data collection strategy that could yield robust subject-specific CAPs when acquisition time is limited and fixed for each participant.

## 2. MATERIALS AND METHODS

### 2.1 Dataset and preprocessing

The publicly available Midnight Scan Club (MSC) dataset, which includes data from ten adults (5 females, age = 29.1 ± 3.3) was used in this work (Gordon, Laumann, Gilmore, et al., 2017). Ten resting-state fMRI scans were collected for each participant on separate days, using a Siemens TRIO 3T MRI scanner with gradient-echo EPI sequence (TR = 2.2 s, TE = 27 ms, flip angle = 90°, voxel size = 4×4×4 mm^3^, 36 slices). Each scan lasts for 30 minutes (818 volumes) and begins at midnight. Subject MSC08 was excluded in this work as the participant fell asleep during the fMRI scanning, hence nine subjects with a total of ninety scans were analyzed in this study. More details about the participant information and MRI acquisition parameters have been described in (Gordon, Laumann, Gilmore, et al., 2017).

The fMRI data of the MSC dataset has been preprocessed and can be accessed through OpenNeuro (https://openneuro.org/datasets/ds000224). In brief, the preprocessing steps include slice timing correction, intensity normalization (mode = 1,000), within-subject head motion correction, and distortion correction. The functional data were registered to the Talairach atlas space using the average T2-weighted image and the average T1-weighted image. To account for anatomical differences between subjects, FSL’s FNIRT was used to non-linearly warp the mean T1 image to the MNI space, and the preprocessed volumetric time series at Talairach space were then registered to the registered T1 image at the MNI space (Greene et al., 2020). Schaefer’s parcellation with 400 cortical regions (Schaefer et al., 2018), which includes the visual network (VN), somatomotor network (SMN), dorsal attention network (DAN), ventral attention network (VAN), limbic network, fronto-parietal network (FPN) and default mode network (DMN) (Yeo et al., 2011), were used to extract the regional BOLD signal. For each ROI, the amplitude of the time series was normalized using z-score independently, and the normalized amplitude at each time point reflects the activity deviation from its own baseline (Liu & Duyn, 2013).

### 2.2 Individual coactivation pattern analysis at the subject- and scan-level

The coactivation patterns can be estimated in a seed-based (Chen et al., 2015; Liu & Duyn, 2013) or seed-free manner (Liu et al., 2013). In accordance with our previous studies, we used a robust pipeline that performs the CAP analysis using a seed-free manner at the ROI-level (Yang et al., 2021; Yang et al., 2022). Besides the group-level analysis, we also constructed the individual CAP states at both subject-level and scan-level (Figure 1A). As for the group-level, the regional time series (818×400) from all the 90 scans were concatenated as the coactivation matrix (73,620×400), where 73,620 (818×90) is the total volumes and 400 is the ROI number. The distance between the whole-brain coactivation profiles (400×1) from any two volumes was calculated using correlation distance (1 – Pearson correlation coefficient), and K-means clustering was then performed to identify brain states with similar coactivation profiles across all volumes. The group-level CAP maps were obtained by averaging volumes assigned to the same state, and normalized by dividing the within-cluster standard deviation (Liu & Duyn, 2013). For the subject-level CAP analysis, ten scans from each participant were grouped as the coactivation matrix (8,180×400), and the same clustering procedure was performed for each subject, respectively. Finally, the scan-level CAPs were generated by applying the CAP pipeline to the coactivation matrix (818× 400) from a single scan of each subject. We estimated the CAP states with different cluster numbers (K = 2, 3… 21), and the silhouette score was calculated to evaluate the clustering results (Figure S1). As we have performed the clustering procedure to obtain CAP states using a single scan, we chose K = 4 as a trade-off to ensure that each cluster has adequate data to generate reliable spatiotemporal dynamics, and the identical state number was used for the group-level and subject-level for easier comparisons.

**Figure 1.**
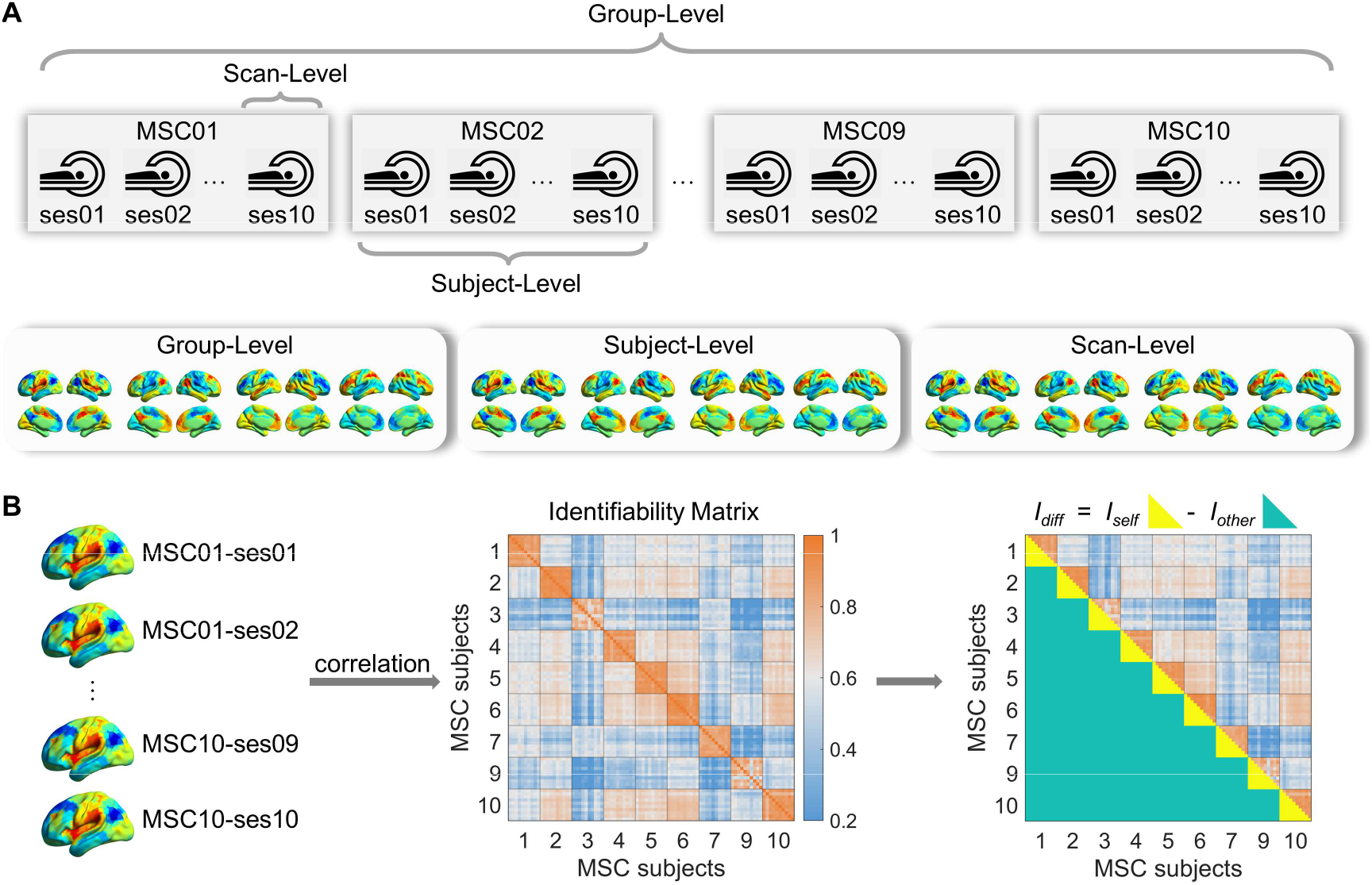
The pipeline for CAP states estimation and subject identification analysis. (A) The CAP analysis was performed at three levels. The group-level CAPs were obtained using scans from all subjects, the subject-level CAPs were generated based on all scans from each individual, and the scan-level CAPs were acquired by using a single scan. (B) The spatial similarity between CAP maps from all the ninety scans was measured by Pearson correlation, and the correlation matrix was defined as the identifiability matrix. Differential identifiability (***I_diff_***) was calculated by subtracting the average intersubject similarity (***I_other_***) from the average intra-subject similarity (***I_self_***).

To match and compare CAPs obtained at different levels in the following analyses, the four group-level CAP states were used as references, and the one-to-one correspondence of CAP pairs between group-level and subject-level/scan-level was established by using the Hungarian algorithm respectively (Gutierrez-Barragan et al., 2019; Kuhn, 1955; Tarun et al., 2020). In addition, after obtaining CAP states at the group-level and subject-level, CAP maps of each scan were further reconstructed by averaging the volumes belonging to every single scan. Therefore, for each of the three levels, there were four CAP states (K = 4) for each single scan, and the following analyses were performed based on reconstructed CAPs from ninety scans at each level independently.

### 2.3 Subject identification and differential identifiability

Recently, Finn and colleagues have shown the uniqueness of functional connectome as a robust brain fingerprinting (Finn et al., 2015). By calculating the Pearson correlation between the sample FC from Day 1 and all FC from Day 2, the other scans from the sample subject exhibited stronger correlations compared with scans from other subjects. In this work, a similar subject identification procedure was performed. First, for each CAP state, the similarity across scans and subjects was measured by calculating the Pearson correlation between the ninety CAP maps (400×90), and the similarity matrix (90×90) was termed as **identifiability matrix** (Figure 1b). As there were ten scans for each subject, the anticipation is that the inter-scan similarity between the ten scans from the same subject should be the highest. Then, for each sample scan, nine scans with the highest similarity were selected from the other 89 scans, and the percentage of correct identifications (i.e., scans selected from the sample subject) was defined as the identification (ID) rate. The CAP variance maps were also generated by calculating between-subject variances of each region based on the linear mixed-effects model. Brain region with a larger variance is supposed to be more important to distinguish between individuals.

Besides the aforementioned target-based subject identification method, another quantification metric, differential identifiability (***I_diff_***) was also used in this study (Amico & Goni, 2018). The ***I_diff_*** is defined as the difference between the average intrasubject similarity (***I_self_***, diagonal of the identifiability matrix) and the average intersubject similarity (***I_other_***, off-diagonal of the identifiability matrix), and a larger ***I_diff_*** indicates better identifiability. All values (***I_self_***, ***I_other_***, ***I_diff_***) were multiplied by 100 for more straightforward results demonstration. To estimate the contribution of a specific region to subject discrimination, the identifiability matrix and differential identifiability were re-calculated by removing the ith ROI (i = 1, 2,…, 400) iteratively, and the changed differential identifiability (***ΔI_diff_***) was obtained by subtracting the original ***I_diff_*** from the new ***I_diff_*** (Jo et al., 2021). A negative **Δ*I_diff_*** means the new ***I_diff_*** decreased after removing the region, hence -**Δ*I_diff_*** was used to indicate the contribution of the ith region. The contribution of each functional network was also estimated using a similar manner by removing all regions from the network and estimating the **Δ*I_diff_***

### 2.4 Effects of principal component reconstruction and head motion

Based on the same dataset (MSC), previous subject identification studies have shown that the ***I_diff_*** can be improved by reconstructing the FC matrix through several primary principal components (PCs) with large explained variance (Amico & Goni, 2018; Jo et al., 2021). We validated this approach in our work, we decomposed the identifiability matrix (90×90) of each CAP state into 89 principal components, and the 89 PCs were sorted in descending order based on their explained variances. Then, the identifiability matrix was reconstructed using the first ***N*** components (***N*** = 1 to 89), and the ***I_diff_*** was re-calculated. Although we validated the improvement of subject identification by using PCA reconstruction (Figure S3), the main results reported in this study were based on the original identifiability matrix without PCA reconstruction.

On the other hand, the detrimental effects of in-scanner head motion have been well-documented before, which could contaminate the BOLD signal and introduce spurious functional connectivity (Power et al., 2012; Van Dijk et al., 2012). In this study, we evaluated how in-scanner head motion affects the CAP state estimation and the subsequent subject identification analysis. The detailed methods, results and discussions about the effects of head motion were described in the supplementary materials.

### 2.5 Explore optimal data collection strategy to estimate robust subject-specific CAP states

The single scan from the MSC dataset used in this study is 30 minutes long, which is longer than most current fMRI studies. It is important to know how much data is required to achieve stable CAP states with sufficient individual differences (high ID rate and large ***I_diff_***). To answer this question, we randomly extracted ***T*** (from 5 to 29) consecutive minute data from each fMRI scan independently. As we found the scan-level CAP states were unstable even using the full data (30 mins per scan), the effects of scan length were only evaluated at the group-level and subject-level. Particularly, with the assumption that *more data is better*, the CAPs obtained by the full data (10 scans per subject, 30 mins per scan) at the group-level and subject-level were used as referenced CAPs. Then, the CAP maps and their differential identifiability were reestimated based on the sampling data, and the spatial similarity between the CAPs estimated by subsamples and the referenced CAPs was measured by Pearson correlation. As the data sampling is random, we repeated the above procedure 100 times, and the average results were reported.

Unsurprisingly, we found more data yielded better results in the aforementioned analysis (Figure 5). However, the acquisition time of each participant is limited in practice, and it is necessary to know what data collection strategy (e.g., more scans with shorter duration, or fewer scans with longer duration) is better to capture reliable individual CAP states. We found that the subject-level CAPs obtained using half of the data (10 scans, 15 mins/scan) were already highly similar to the referenced CAPs generated by full data (Figure 6). Therefore, we equally sampled a total of 150 minutes of data from 5 (30 mins/scan) to 10 scans (15 mins/scan) for each participant, estimated the subject-level CAPs, and measured their spatial similarity with the referenced CAPs. While one potential issue that exists for this approach is that the referenced CAPs were obtained using data from all scans, hence the higher similarity achieved by more scans with shorter duration might be caused by information leakage. To avoid this problem, we performed another experiment by randomly selecting two non-overlapped subsets with fixed data length (60 minutes) for each subject, and the scan number (*N*) ranged from *N* = 2 (30 mins/scan) to *N* = 5 (12 mins/scan). The subject-level CAPs were then estimated in each subset, and the spatial similarity between the two subsets was compared by Pearson correlation.

## 3. RESULTS

### 3.1 Coactivation patterns generated at distinct levels

Four CAP states were identified and analyzed at the group-, subject- and scan-level independently, and CAP maps obtained at the subject-level and scan-level were aligned to the group-level CAPs. The subject-level and scan-level results were averaged across subjects and scans, and are presented in Figure 2, and the coactivation level of each functional network is shown in Figure S2. Two CAP pairs with opposite coactivation profiles were obtained. For instance, State 1 was mainly dominated by the activated VAN, SMN and deactivated DMN, and the opposite pattern was observed in State 2. Similarly, State 3 showed stronger deactivation in the FPN, DAN and activation in the anterior DMN, and the opposite for State 4. It can be observed that the degree of co-activation or co-deactivation level (Z-value) of each functional network decreased from group-level to scan-level, suggesting the underlying inter-subject and intra-subject variations across subjects and scans.

**Figure 2.**
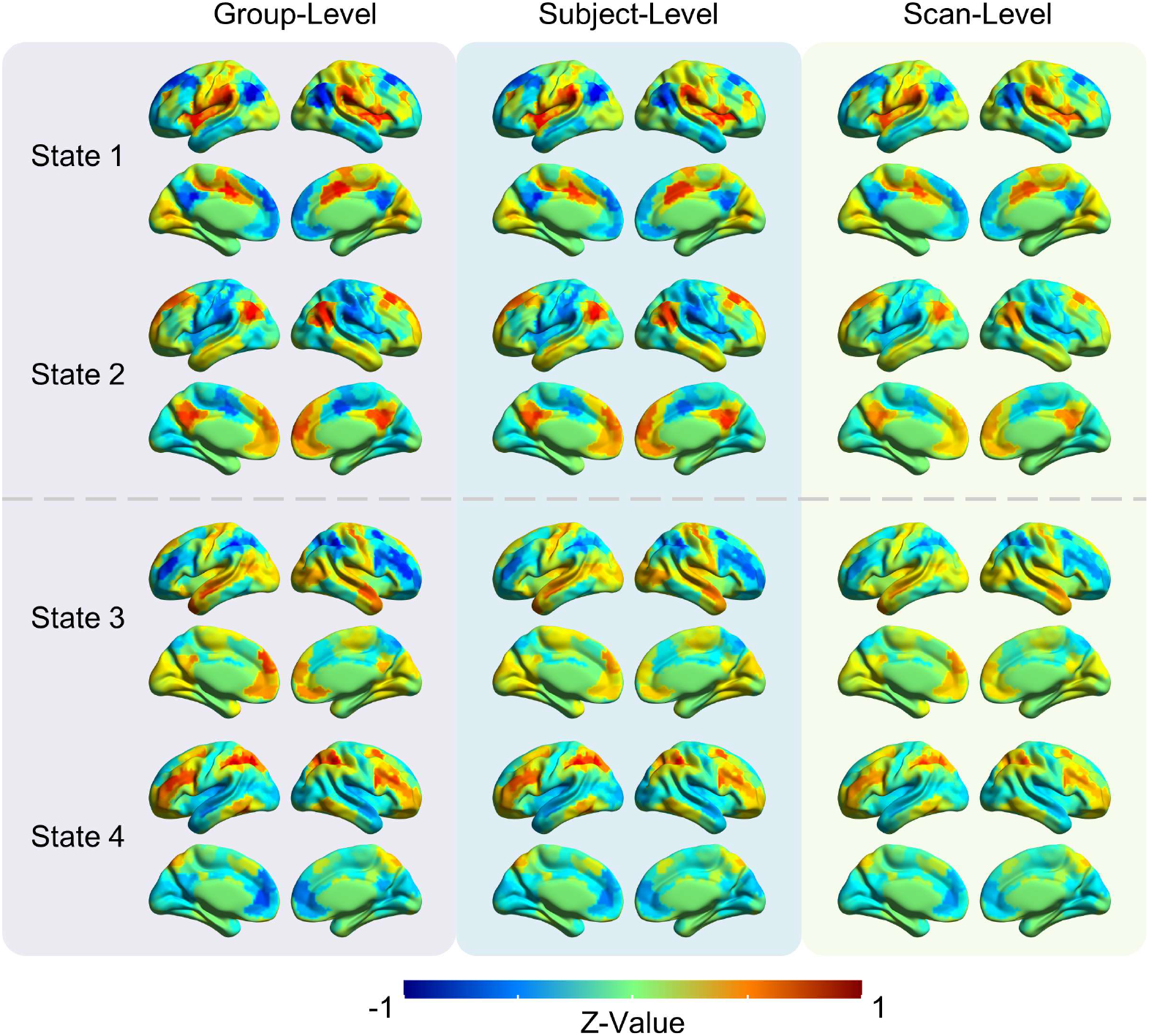
The group-averaged CAP maps obtained at the group-level, subject-level and scan-level. Four CAPs were obtained in this work, and two CAP pairs with opposite spatial patterns were identified. State 1 and 2 were mainly dominated by stronger activation or deactivation in the DMN, VAN and SMN. State 3 and 4 were primarily dominated by stronger activation or deactivation in the FPN, DAN and anterior DMN. The red color indicates a stronger activation level than the baseline of that region, and vice versa for the blue color.

### 3.2 Individual CAPs at the subject-level achieved the best identifiability

Based on the identifiability matrix, we calculated the identification (ID) rate and differential identifiability (***I_diff_***) to evaluate the subject identification performance of each CAP state at different levels. As shown in Figure 3A, we can identify two opposite CAP pairs from the identifiability matrix. Within each state, scans from the same subject were more similar than scans from different subjects, and the identifiability matrix of State 1 was further extracted as a finer illustration (Figure 3B). As reflected by the diagonal elements of the identifiability matrix, both the group-level and subject-level CAPs exhibited high intra-subject similarity. However, only MSC02 and MSC06 showed consistently high intra-subject similarity across the ten scans at the scan-level. On the other hand, compared with the group-level, lower inter-subject similarity can be observed at the subject-level from the off-diagonal values.

**Figure 3.**
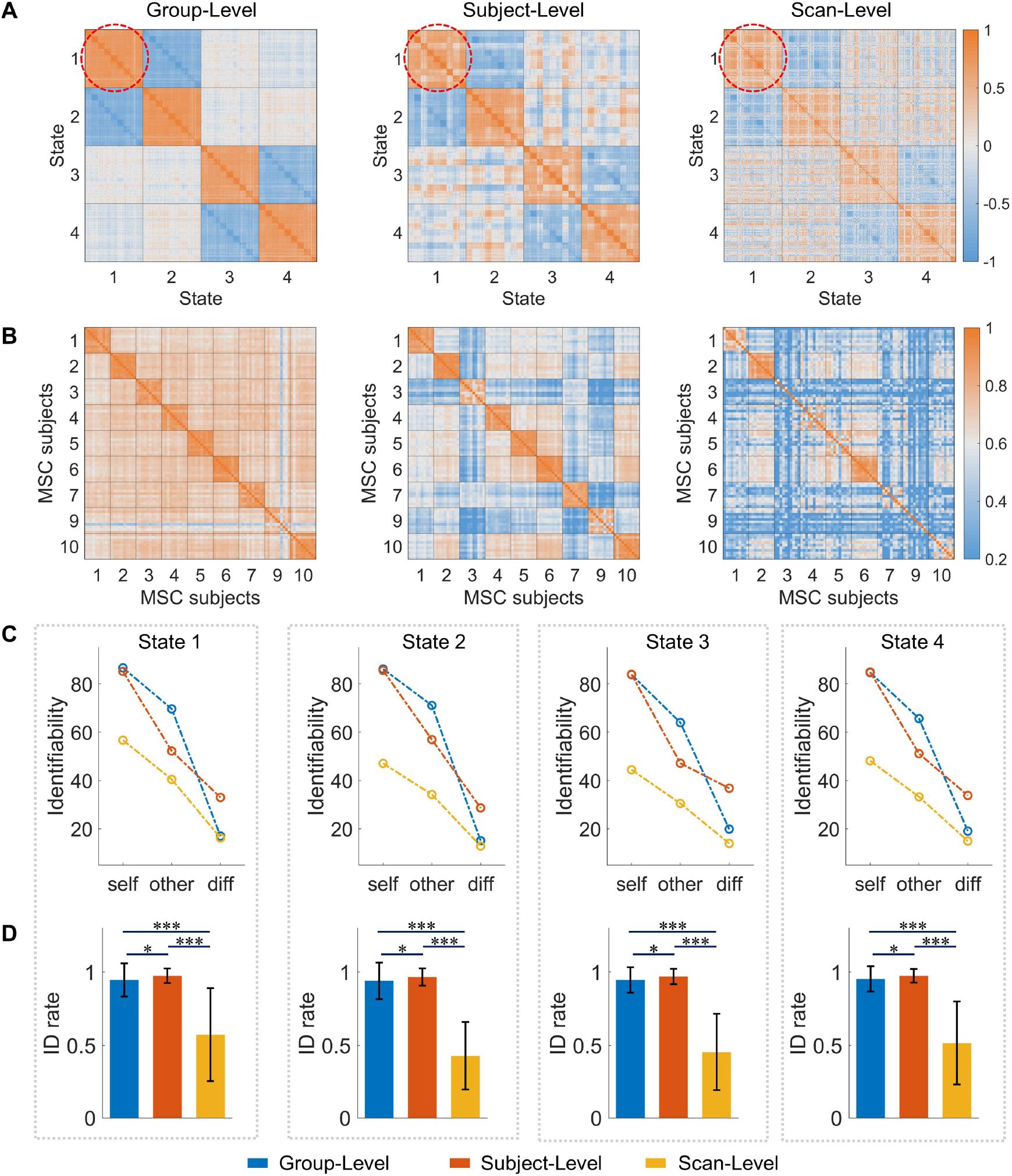
Identifiability matrices and subject identification performances at the three levels. (A) Identifiability matrix of the four CAP states. (B) Identifiability matrix of State 1. The colorbar shows the Pearson correlation coefficient. (C) The intra-subject similarity (***I_self_***), inter-subject similarity (***I_other_***), differential identifiability (***I_diff_***), and (D) the identification (ID) rate at the three levels. Permutation test was performed between different levels to compare their differences in ID rate, and false discovery rate (FDR) was used to correct multiple comparisons. * indicates P < 0.05, and *** indicates P < 0.0001. Errorbar shows the standard deviation.

The identification accuracy and differential identifiability were then calculated based on the identifiability matrix from each state separately. As shown in Figure 3C, both high intra-subject similarity (***I_self_*** > 80) and inter-subject similarity (***Iother*** > 60) across the four CAP states were obtained at the group-level. On the contrary, scan-level CAPs were characterized by low ***I_self_*** (< 60) and ***I_other_*** (< 40). Hence, group-level and scan-level exhibited low differential identifiability (***I_diff_*** < 20). However, as shown in Figure 3D, the ID rate achieved by the group-level (> 90%) was much higher than the scan-level (< 50%). As for subject-level CAPs, while maintaining high intra-subject similarity, the inter-subject differences were also enlarged, which leads to the highest differential identifiability among the three levels. Besides, subject-level also achieved the highest ID rate. The detailed values of the ID rate and ***I_diff_*** can be found in the supplementary Table S1 and S2.

In addition, we also used PCA to decompose the identifiability matrix, and reconstructed the identifiability matrix by the linear combination of different numbers of PCs. The optimal ***I_diff_*** values at the three levels were listed in the supplementary Table S3. In brief, PCA-optimized CAP states showed increased ***I_diff_*** across all levels, and the subject-level CAPs achieved the maximum growth. Particularly, the results of State 3 at the subject-level are shown in Figure S3 as an example. The ***I_diff_*** increased from 36.77 to 42.59 when the identifiability matrix was reconstructed using 9 PCs.

Finally, we evaluated how in-scanner head motion affects the estimation of CAP states and subject identification. We observed that participants with higher intra-subject similarity (e.g., MSC-06) and CAP states with larger ***I_diff_*** (e.g., State 3) contained fewer motion-contaminated frames (Figure S7). Particularly, the quantitative analyses suggest that motion-contaminated frames would reduce the intra-subject similarity, which resulted in a lower ID rate and ***I_diff_*** (Figure S8). Additionally, the ***I_diff_*** increased a little bit after excluding the motion-contaminated frames (Figure S9 and Table S4). More detailed results and discussions have been described in the supplementary materials.

### 3.3 Higher-order cortical regions contributed to the subject identification

To investigate the contribution of each brain region to subject identification, intersubject variance maps were calculated for each CAP state, and we mainly reported the subject-level results in the main text. As shown in Figure 4A, opposite CAP pairs showed similar variance patterns across the brain. For State 1 and 2, brain regions with larger variances were mainly located in the left hemisphere, including the middle frontal gyrus, superior parietal lobe and supramarginal gyrus. For State 3 and 4, the bilateral medial and superior prefrontal cortex, anterior cingulate cortex, postcentral gyrus and the right superior parietal lobe showed larger variances.

**Figure 4.**
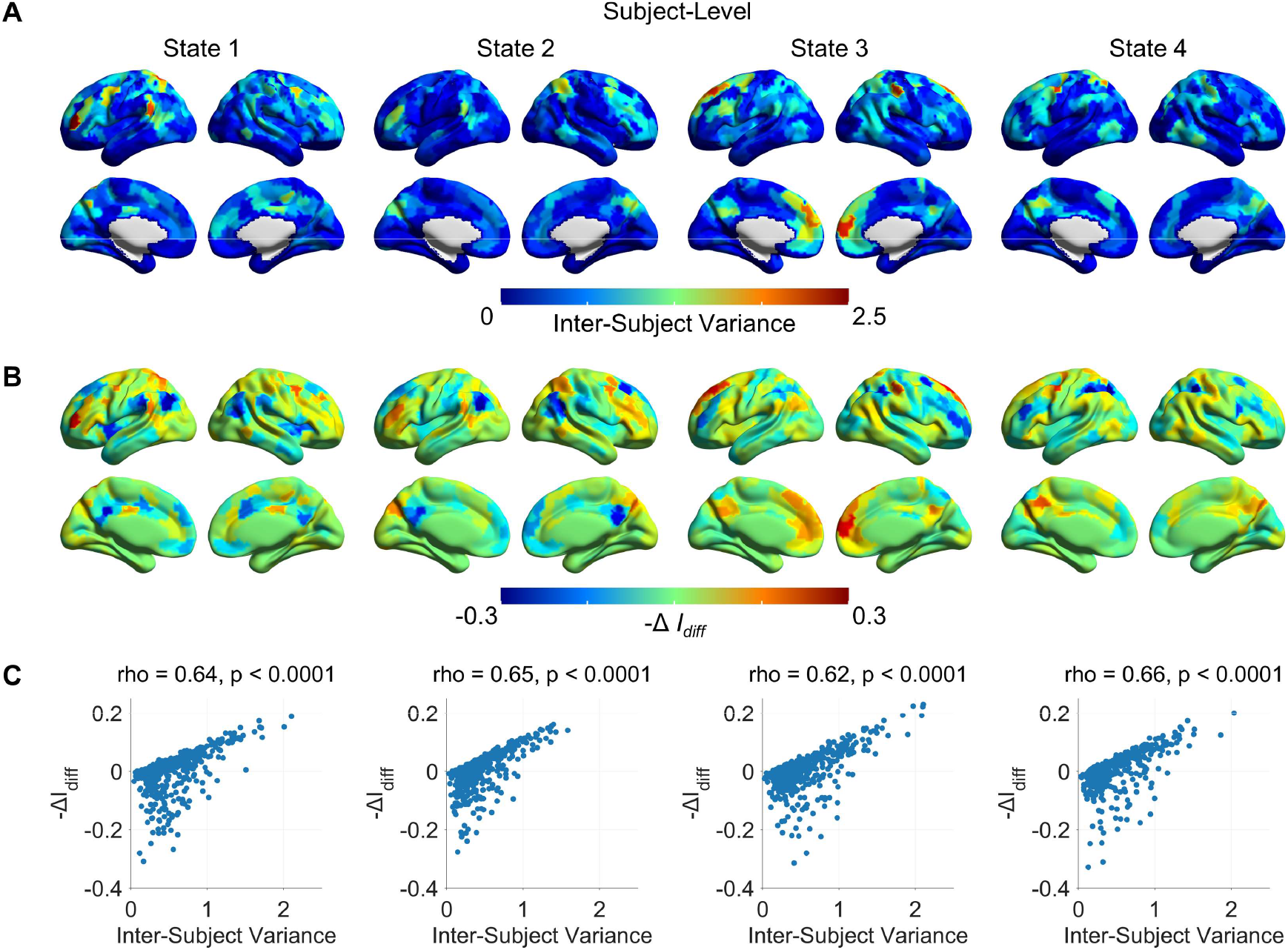
Contribution of brain regions to the subject identification at the subject-level. (A) The inter-subject variance maps. (B) The changed differential identifiability (-**Δ*I_diff_***) maps. (C) A significant positive correlation was found between the inter-subject variance map and the -**Δ*I_diff_*** map for each CAP state.

In addition, we also measured the changed differential identifiability (**Δ*I_diff_***) of each brain region. The -**Δ*I_diff_*** was reported because a positive -**Δ*I_diff_*** indicates a greater contribution of the region to the individual differences. Similar to the results of variance, -**Δ*I_diff_*** maps varied between CAP pairs. For example, brain regions belonging to the FPN and DAN showed a larger -**Δ*I_diff_*** in State 1 and 2, while the DMN-related regions showed a larger -**Δ*I_diff_*** in State 3 and 4. Finally, we compared the pattern of the variance map and -**Δ*I_diff_*** map by calculating Spearman’s rank correlation, and a significant positive correlation was found for each CAP state (rho > 0.6, p < 0.0001).

As for the variance and -**Δ*I_diff_*** maps at the group-level and scan-level, similar distributions with lower regional contribution values were found, see Figure S4 and Figure S5. Furthermore, we measured the contribution of each functional network by excluding regions belonging to each network. As shown in Figure S6, similar -**Δ*I_diff_*** maps were obtained at the three levels, and networks at the subject-level exhibited overall larger -**Δ*I_diff_*** values than the other two levels.

### 3.5 Longer scan length generated more reliable subject-specific CAP states

As the scan length of a single resting-state fMRI run used in the current study is 30 mins, we tested how scan length (from 5 to 30 mins) would affect the CAP estimation and subject identification. In summary, as shown in Figure 5, the subject identification performance (ID rate and ***I_diff_***) increased with the scan length at both the group-level and subject-level, and the spatial similarity to the referenced CAPs (obtained using 30 mins fMRI data) also increased with the scan length. Particularly, these values increased rapidly from 5 to 10 mins, and gradually slowed down from 10 to 15 mins. It can be observed that the performance achieved by 15-min data per scan (in total 150 mins per subject) is comparable with the results of 30-min data per scan (in total 300 mins per subject). For instance, for the four CAPs at the subject-level, the ID rates at 15 mins were around 95%, and their spatial similarities to the referenced CAP were above 0.95.

**Figure 5.**
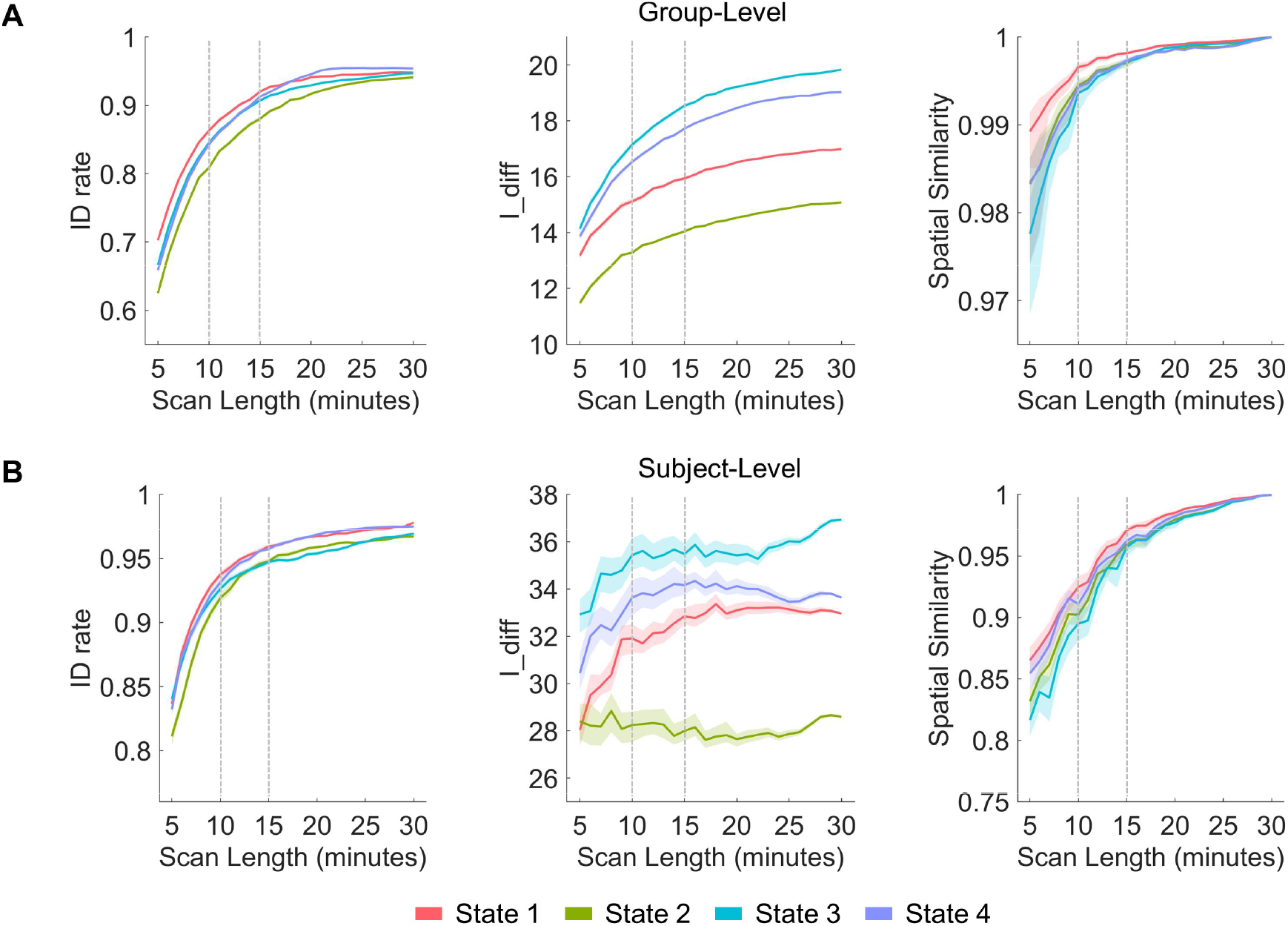
The effects of scan length at the (A) group-level and (B) subject-level. The first two columns show the ID rate and ***I_diff_***, and the last column shows the spatial similarity to the CAPs obtained using 30 mins of fMRI data. Two vertical dashed lines indicate the results at 10 and 15 mins. The shadow is the 95% confidence interval of the mean.

Next, we tested how different data collection strategy affects the estimation of subject-specific CAPs with a fixed amount of data. As we found that CAP states obtained using 150-min data were highly similar to the referenced CAPs (300-min data), we re-estimated the subject-level CAPs using 150-min data with different strategies, from 5 scans (30 mins/scan) to 10 scans (15 mins/scan), and compared them with the referenced CAPs. As shown in Figure 6, the results suggested that the CAPs estimated by 5 scans (30 mins/scan) showed the lowest spatial similarity. More scans with shorter duration generated more similar subject-level CAPs compared with the referenced CAPs for all four states, and reached the maximum with 10 scans and 15 mins/scan.

**Figure 6.**
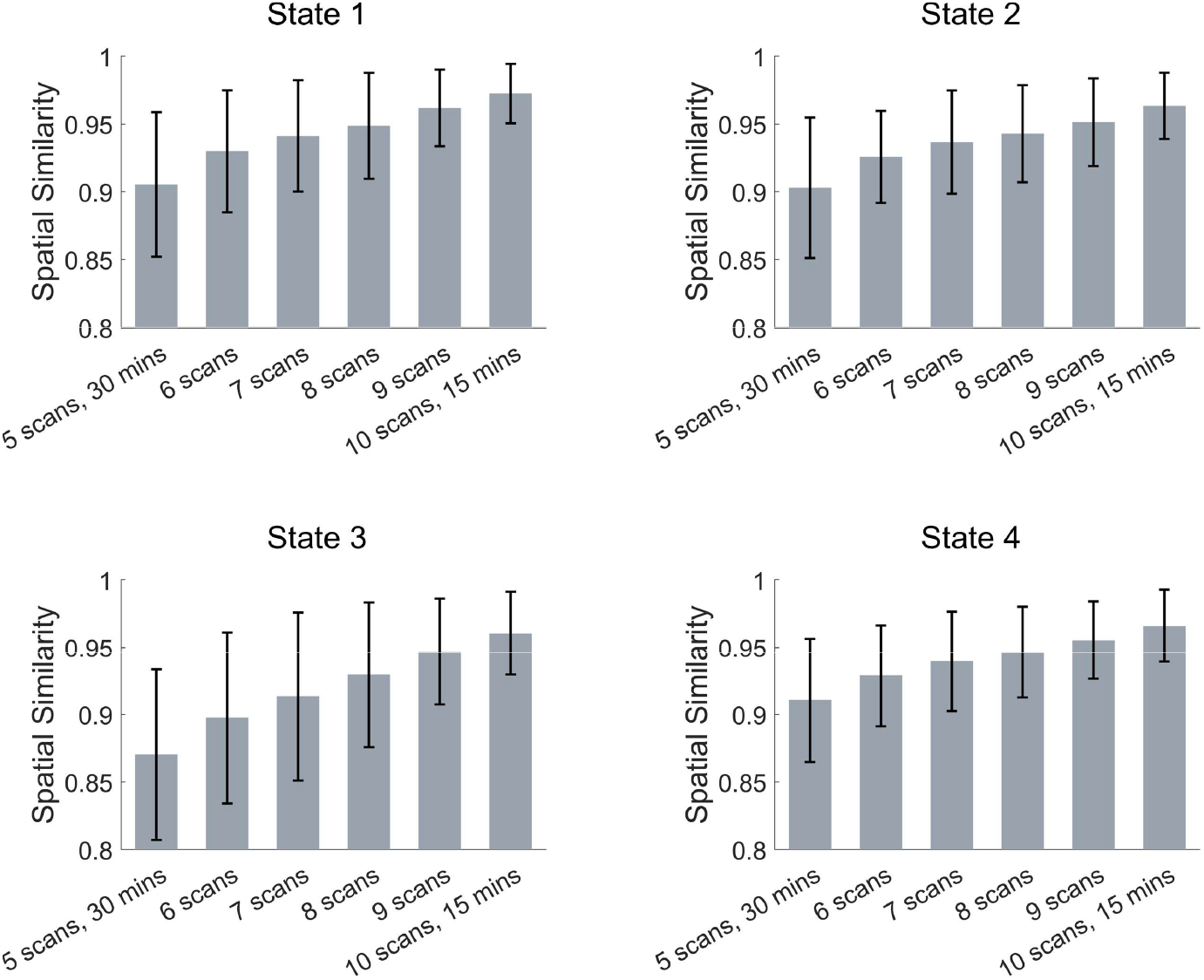
The spatial similarity between subject-level CAPs estimated by a fixed length of data and the referenced CAPs. The 150-min data were equally sampled from 5 (30 mins/scan) to 10 scans (15 mins/scan), and the random sampling was performed 100 times. With the same total length, CAP states estimated by more scans but shorter duration were more similar to the referenced CAPs generated by all data (10 scans/subject, 30 mins/scan). The spatial similarity was measured by Pearson correlation, and the errorbar shows the standard deviation.

However, the increased spatial similarity achieved by more scans and shorter duration could be caused by the information leakage, as the referenced CAPs were obtained using all scans. Therefore, instead of comparing with the referenced CAPs, we randomly selected two non-overlapped subsets with the same scan number (*N* = 2 to 5) and fixed total length (60 mins) for each participant, and compared the subject-level CAPs between the two subsets (Figure 7). It can be observed that subject-specific CAPs were less similar between the two non-overlapped subsets when only 2 scans (30 mins/scan) were included in each subset. For State 1 and 3, the subject-specific CAPs showed the highest inter-subset similarity with 4 scans (15 mins/scan) within each subset, and the highest inter-subset similarity for State 2 and 4 were obtained using 5 scans (12 mins/scan) within each subset.

**Figure 7.**
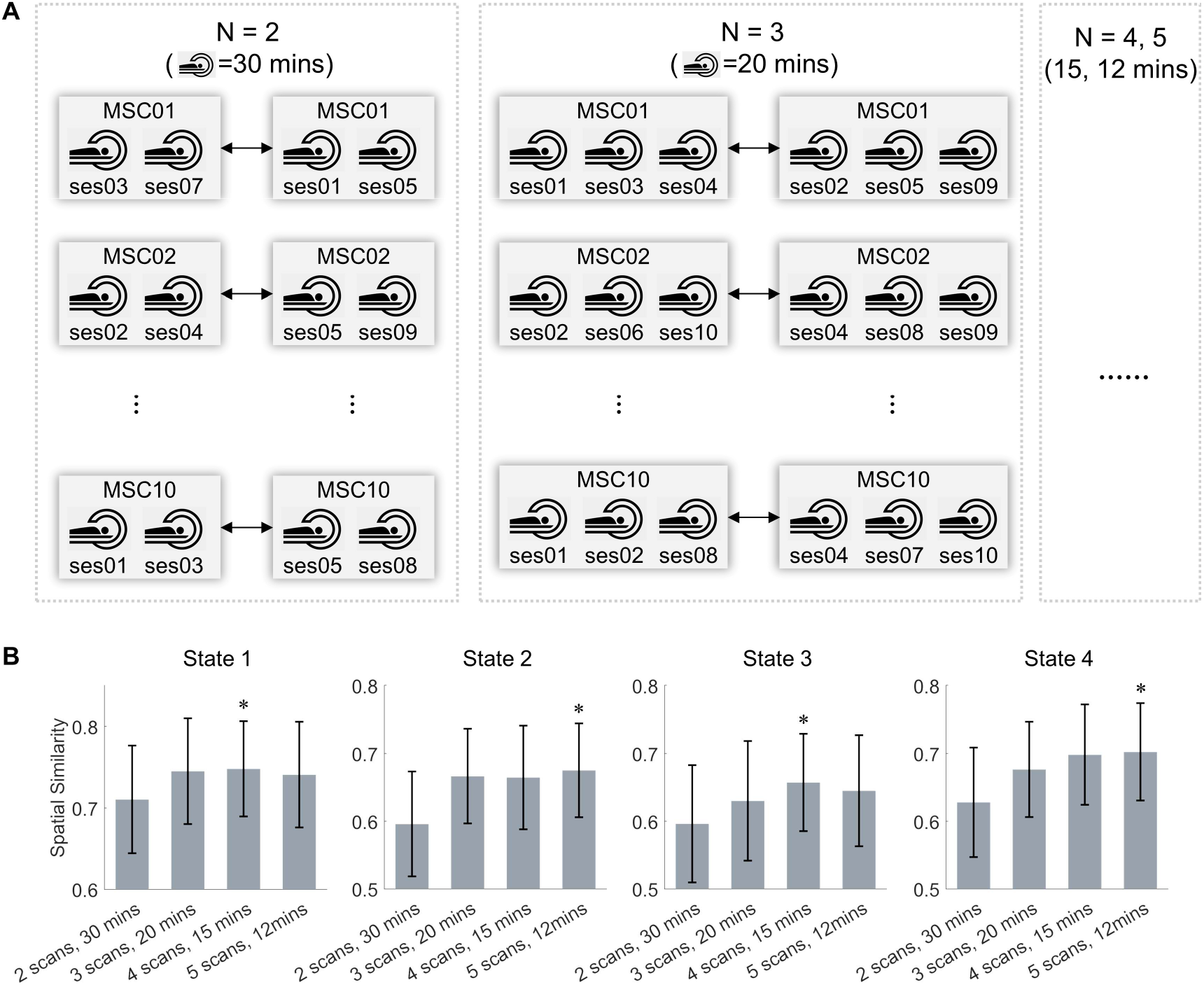
The spatial similarity between subject-level CAPs from two non-overlapped subsets. (A) For each subject, two non-overlapped subsets with equal scan numbers (*N* = 2 to 5) were randomly extracted from the ten scans, and the subject-level CAPs were estimated for the two subsets independently and compared. Random sampling was performed 100 times. (B) The intra-subject spatial similarity for the four CAP states between two subsets. The intra-subject similarity increased when more scans were included in each subset. The spatial similarity was measured by Pearson correlation, and the errorbar shows the standard deviation. The largest value was labeled by *.

Finally, we further validated the notion that *more data is better* by calculating inter-subset similarity for each participant with an increased amount of data (i.e., using different scans (*N* = 2 to 5) with the same duration (30 mins/scan)). As shown in Figure S10, the inter-subset similarity increased with more scans used in each subset, suggesting the subject-specific CAPs are more generalizable when more scans are used.

## 4. DISCUSSION

The present study went beyond the commonly used group-level analysis by utilizing individual participant data to estimate CAP states at both the subject-level and scan-level. Our findings revealed that enlarging inter-subject differences while preserving high intra-subject similarity led to the highest identification accuracy and the largest differential identifiability of individual CAPs at the subject-level. The identification process was primarily facilitated by brain regions within the association networks, such as the FPN, DAN, and DMN. Conversely, obtaining reliable individual CAPs through a single prolonged scan is a challenging task. In the end, we validated that more data is better for CAP estimation and subject identification, and 150-min data per subject is enough to obtain robust subject-specific CAPs with sufficient individual differences. We also recommend collecting more scans with shorter duration (12~15 mins/scan) given a fixed and limited acquisition time in practice.

### 4.1. Subject-level CAPs enlarged the inter-subject discrimination and improved the subject identification accuracy

In this study, four CAPs were generated at the group-level, subject-level and scan-level, respectively. Due to the low intra- and inter-subject similarity, the scan-level failed to identify subjects accurately, and its differential identifiability was low. As for the group-level, the high intra-subject similarity resulted in a high ID rate, while it also exhibited high inter-subject similarity, which also led to the low ***I_diff_***. However, the subject-level CAPs maintained the high intra-subject similarity and enhanced the intersubject differences, hence the highest ID rate and largest ***I_diff_*** were obtained at the subject-level. Similarly, a recent edge-centric FC study found that, by clustering high-amplitude cofluctuation events, the co-fluctuation states estimated at subject-level were personalized and could predict the subject’s FC better than group-averaged states, which highlighted the importance of individualized brain features (Betzel et al., 2022).

Overall, the group-level and subject-level achieved comparable identification rate with previous studies (Finn et al., 2015; Menon & Krishnamurthy, 2019). As for the differential identifiability, previous node-FC studies have found that ***I_diff_*** could be optimized by using PCA reconstruction (Amico & Goni, 2018; Bari et al., 2019). With the benefits of higher-order reconstruction, edge-centric FC further improved the subject idiosyncrasies (Jo et al., 2021). In the current study, group-level and scan-level CAPs showed comparable differential identifiability (***I_diff_*** < 20) to previous node-FC results (Amico & Goni, 2018; Bari et al., 2019). Consistent with previous studies (Amico & Goni, 2018; Bari et al., 2019; Jo et al., 2021), PCA-optimized reconstruction further improved the differential identifiability. The PCA-optimized ***I_diff_*** values of three CAP states (from 38.52 to 42.59) at the subject-level were better than the best results achieved in the same dataset (MSC) before (***I_diff_*** = 35.27, based on edge-centric FC, PCA optimized) (Jo et al., 2021). Notably, even the original ***I_diff_*** at the subject-level of State 3 (***I_diff_*** = 36.77) was slightly higher than the PCA-optimized ***I_diff_*** of edge-centric FC (***I_diff_*** = 35.27), suggesting that despite the fact that coactivation profiles are first-order brain measurements, brain dynamics could provide additional personal traits and improve the subject identification (Menon & Krishnamurthy, 2019).

### 4.2. Brain regions of association networks were more discriminative

To evaluate the impact of brain regions and networks on subject identification, we calculated the change in differential identifiability by removing one region or network at a time (Jo et al., 2021). The most discriminative brain regions were mainly located in the higher-order networks, such as the FPN, DAN and DMN. These higher-order association cortices are critical for cognition (Cole et al., 2013), the most evolutionarily recent (Sepulcre et al., 2010), and characterized by large individual variability (Laumann et al., 2015; Mueller et al., 2013). Together, these might be the reasons why these heteromodal regions drive the subject identification (Finn et al., 2015; Horien et al., 2019; Jo et al., 2021). In addition to the changed differential identifiability (-**Δ*I_diff_***), the current study also calculated the inter-subject variance of CAPs. Strong positive correlations were found between the variance maps and -**Δ*I_diff_*** maps, suggesting that brain regions with larger individual differences tend to be more identifiable. Moreover, the distribution of the regional contribution (-**Δ*I_diff_***) varied among CAP states, indicating that different information was encoded in distinct dynamic brain states.

### 4.3. Individual CAP states were unstable over time at the scan-level

An important question is whether stable individual CAP states could be identified in acquisitions over time. For scan-level CAP analysis, although the length of a single scan is long (30 mins/scan), consistent CAPs across the ten scans were only obtained in a few subjects (e.g., MSC02, MSC06), which is different from previous studies that demonstrate individual connectome is stable over time or cognitive states (Gratton et al., 2018; Horien et al., 2019; Seitzman et al., 2019). This could be attributed to the fact that the previous individual connectome was based on static FC, which remains relatively stable across both resting-state and task conditions as an intrinsic functional architecture (Cole et al., 2014). However, the reliability of dynamic FC was found to be lower than that of static FC, with more dynamic connections being less reliable (Zhang et al., 2018). Although we have demonstrated that the group-level CAPs are reproducible across different analytic approaches and can be generalized to independent datasets (Yang et al., 2021), the test-retest reliability of individual CAPs across multiple scans for the same participant remains to be investigated. Besides, one previous study found that the brain state could be manipulated by scan conditions (Finn et al., 2017), and the state-specific individual functional parcellation is not feasible with a single acquisition (Salehi, Greene, et al., 2020). As the underlying unrestricted and unobservable mental process could be varied across resting-state sessions for the same subject, the brain might reconfigure at distinct cognitive states (Krienen et al., 2014; Salehi, Karbasi, et al., 2020) and cause the unstable individual CAP states when using a single scan. Consequently, multiple scans are necessary to estimate robust individual CAPs at the subject-level.

### 4.4. Collecting more scans with shorter duration improved the reliability of individual CAP states

Previous static FC-based subject identification studies have investigated the effect of scan length on subject identification, and found a longer scan length led to a higher identification accuracy and larger differential identifiability (Amico & Goni, 2018; Bari et al., 2019; Jo et al., 2021). Our findings are in line with these studies and suggest that longer scan length improved subject identification, as indicated by higher ID rate and larger ***I_diff_***. Based on the idea that *more data is better*, CAP states obtained by the full data (10 scans per subject, and 30 mins/scan) were defined as referenced CAPs, and CAPs estimated by longer duration exhibited higher spatial similarity to the referenced CAPs. Particularly, subject-specific CAPs derived from half of the data (10 scans per subject, and 15 mins/scan) were already highly similar (r > 0.95) to the referenced CAPs, and have achieved high identification accuracy and large differential identifiability. Thus, based on our findings, it is recommended to collect a total of two and a half hours of data for each participant to obtain reliable subject-specific dynamic brain states.

Although, sufficient temporal information (i.e., time points) is required to obtain CAP states with robust spatial profiles and accurate temporal dynamics (Liu et al., 2018). While in practice, the total acquisition time of fMRI is not infinite, and the participant could feel uncomfortable and thus produce more in-scanner head motion if a single scan is too long (Meissner et al., 2020). Therefore, in the case of limited total scan length (e.g., 150-min data per subject), it is important to implement an appropriate approach for data collection. Our results revealed that collecting more scans with shorter duration (e.g., 10 scans per subject, and 15 mins per scan) is a better approach than collecting fewer scans with longer duration (e.g., 5 scans per subject, and 30 mins per scan). This discovery aligns with prior studies that revealed higher reliability could be attained by employing multiple scans with shorter durations as opposed to a single long scan (Cho et al., 2021; Laumann et al., 2015). The advantages of acquiring more scans with shorter durations may be attributed to several factors. For instance, the impacts of noises (e.g., scanner noise) are more likely to be mitigated through averaging across multiple runs (Akhrif et al., 2018), and individual fMRI data with fewer scans are more susceptible to intra-individual variances. Although the strategy of collecting more scans with shorter duration may have resulted in an overestimation of the similarity to referenced CAPs due to potential information leakage, the validation of subject-specific CAPs from two non-overlapped subsets of the same participant supports the notion that this strategy improves generalizability.

### 4.5. Limitations and future directions

The primary focus of the present study was on the spatial patterns of subject-level CAP states, and further investigation is required to determine if their temporal dynamics are personalized and linked to individual behaviors. Moreover, due to the small sample size (n=10) in the MSC dataset, a larger sample size would be necessary for future studies (Marek et al., 2022). A recent study by Peng and colleagues defined prior CAP states at the group level using a large population and applied them to new subjects. They found that the occurrence rate of individual CAPs was reliable and related to intersubject variability (Peng et al., 2023). Nevertheless, the temporal dynamics of subject-specific CAPs derived from an individual’s own data have not been fully explored, and there is a need to investigate other temporal features, such as dwell time and state transitions, in addition to occurrence rates. Besides, we have assessed the impact of head motion on our results from several aspects (Figure S7-S9), while the effects of different head motion control strategies in the preprocessing pipeline have not been evaluated in this work as we utilized preprocessed fMRI images from the OpenNeuro, and future studies should take different head motion control strategies into account. In addition to resting-state fMRI, recent research has demonstrated that naturalistic stimuli, such as movie watching, can elicit rich and reliable brain state dynamics (Meer et al., 2020) and result in more precise subject identification (Vanderwal et al., 2017). How the spatiotemporal dynamics of coactivation patterns behave under the naturalistic stimulus, and whether it could improve the individual idiosyncrasies can be explored by future studies. Finally, rather than using the group-level atlas, personalized parcellations should also be considered to evaluate the individual CAPs.

## 5. CONCLUSIONS

Individualized CAP states can be reliably obtained at the subject-level rather than the scan-level. Particularly, subject-level CAPs improved the subject identification by capturing both high intra-subject similarity and large inter-subject differences, and heteromodal regions from the higher-order networks mainly contributed to the identifiability. In addition, we recommend collecting at least 150 minutes of data for each participant with appropriate scan duration (~15 mins/scan) to achieve stable individual CAPs and high identifiability ability.

## DATA AVAILABILITY

The preprocessed Midnight Scan Club (MSC) data used in this study are available in the OpenfMRI data repository at https://openneuro.org/datasets/ds000224.

The code used for CAP state can be found in https://github.com/davidyoung1994/CoactivationPattern.

## ETHICS APPROVAL AND PATIENT CONSENT

The study was approved by the Washington University School of Medicine Human Studies Committee and Institutional Review Board. The written informed consent was obtained from all participants.

## ACKNOWLEDGEMENTS

We thank Evan M. Gordon and his team for collecting and sharing the midnight scan club data. This work was supported by the National Natural Science Foundation of China (NSFC) grant (No. 61871420 and 62071109).

## AUTHOR CONTRIBUTIONS

**Hang Yang**: Conceptualization, Methodology, Software, Data Analysis, Writing Original Draft, and Editing. **Xing Yao**: Data Analysis, and Reviewing. **Hong Zhang**: Data Analysis, and Reviewing. **Chun Meng**: Reviewing, Methodology and Editing. **Bharat Biswal**: Conceptualization, Methodology, Software, and Editing.

