## Supplementary Materials for "Estimating dynamic individual coactivation patterns based on densely sampled resting-state fMRI data and utilizing it for better subject identification"

### Methods

#### Effects of head motion on CAP estimation and subject identification

This study evaluated the effects of head motion on CAP state construction and subject identification. First, the framewise displacement (FD) was calculated to measure the head motion (Power et al. 2012), and a temporal mask of each scan was created by labeling time points with FD > 0.2 mm as motion-contaminated frames. The number of motion-contaminated frames was used to represent the head motion level. We first summarized the head motion for each subject, and then the head motion for each CAP state at the group-, subject- and scan-level (Figure S7). Particularly, scans with more motion-contaminated frames are supposed to be less similar to the other scans from the same subject. Therefore, the spatial similarity of $\boldsymbol{scan_i}$ from $\boldsymbol{subject_j}$ to the other nine scans from the same $\boldsymbol{subject_j}$ was averaged and defined as $\boldsymbol{I_{self}(i, j)}$. Then, Spearman correlation was calculated between the head motion level and $\boldsymbol{I_{self}(i, j)}$ across the ninety scans. In addition, the ten scans of each subject were ordered based on the number of motion-contaminated frames, and the top/bottom five scans of each subject were assigned to the high/low motion group, respectively. Then, the two groups' identification rates were compared using a permutation test (5,000 times).

Furthermore, we explored how the inclusion and exclusion of motion-contaminated frames affect the estimation of CAP states and their identification ability. First, we re-estimated the CAP states based on low-motion frames (FD < 0.2 mm) at each level, and compared the spatial similarity between CAPs with and without motion-contaminated frames by Pearson correlation. Besides, for group-level and subject-level, the reconstructed CAPs of each scan were also compared before and after excluding motion-contaminated frames. Then, the differential identifiability ($\boldsymbol{I_{diff}}$) was compared between motion-included CAPs and motion-excluded CAPs.

### Results

**The effects of head motion on CAP estimation and subject identification**

Head motion is often considered as a confounding factor in fMRI studies. With the visual inspection of Figure S7A and Figure S7B, we found that several scans showed much lower intra-subject similarity than other scans from the same subject, and these scans included more motion-contaminated (FD > 0.2 mm) frames. Therefore, we estimated the effects of head motion level on intra-subject similarity for each CAP state, by calculating the Spearman's rank correlation between the number of motion-contaminated frames of each scan and the $\boldsymbol{I_{self}(i, j)}$ separately. Not surprisingly, a significant negative correlation was found for each CAP state, suggesting the scan with more motion-contaminated frames was less similar to scans from the same subject (Figure S8). The scatter plots of State 1 are presented in Figure S8B as examples. We also compared the ID rate between the high-motion group (top five scans per subject) and the low-motion group (bottom five scans per subject). Generally, the low-motion group exhibited significantly higher ID rates across the four CAP states (Figure S8C).

The above results were derived from the fMRI data with motion-contaminated frames. The CAP states were re-estimated at the three levels using data excluding motion-contaminated frames and compared with motion-included CAPs. As shown in Figure S9 and Table S4, the spatial patterns of group-level CAPs and subject-level CAPs remained highly similar before and after the exclusion of motion-contaminated frames, and their $\boldsymbol{I_{diff}}$ values were also comparable. Therefore, for a dataset with vast amounts of data per subject, the spatial patterns of group-level and subject-level CAPs were relatively robust to head motion. As for the reconstructed CAPs for each scan (Figure S9C), compared with the results of group-level and subject-level, scan-level were more affected by motion-contaminated frames. Particularly, after excluding motion-contaminated frames, the CAPs of several scans were negatively correlated with the CAPs obtained using motion-included data. On the other hand, scan-level also achieved the largest increment in $\boldsymbol{I_{diff}}$ after excluding motion-contaminated frames. These results suggest that the scan-level CAPs are more sensitive to head motion.

**Discussion**

**Head motion reduced the intra-subject similarity and identification accuracy**

Head motion is a vital nuisance factor controlled in fMRI studies (Power et al. 2014; Power et al. 2018). In-scanner head movement could produce brain structure artifacts (Zaitsev, Maclaren, and Herbst 2015), contaminate the BOLD signal, and introduce spurious functional connectivity (Power et al. 2012; Van Dijk, Sabuncu, and Buckner 2012). However, besides the side effects to neuroimaging, the head motion has shown homogeneous and trait-like properties (Couvy-Duchesne et al. 2014; Siegel et al. 2017), and exhibited high test-retest reliability (Zuo and Xing 2014). Furthermore, subject-specific spatiotemporal signatures could be induced by head motion (Bolton et al. 2020; Xifra-Porxas, Kassinopoulos, and Mitsis 2021). These findings seem to imply that the head motion might contribute to the subject identification. However, previous studies reported that, compared with functional connectivity profiles, head motion only achieved a much lower above-chance level in identification rate (Finn et al. 2015; Vanderwal et al. 2017).

Consistent with previous findings (Horien et al. 2019), we found that scans with more head motion showed lower inter-scan similarity (Figure S8). Besides, the CAP state with the largest differential identifiability ($I_{diff}$ = 36.77, State 3) was also less contaminated by head motion (Figure S7D), indicating that head motion would reduce the subject identifiability (Amico and Goni 2018). Previous studies found that low-motion subjects achieved a higher identification rate than using all subjects (Horien et al. 2019), and controlling the effects of head motion during the preprocessing increased the accuracy (Xifra-Porxas, Kassinopoulos, and Mitsis 2021). The current study also found that the low-motion group showed a significantly higher identification rate than the high-motion group. Furthermore, the CAP maps and $I_{diff}$ were re-estimated by excluding motion-contaminated frames. The overall spatial patterns remained unchanged for group-level and subject-level CAPs, and the $I_{diff}$ increased for most cases (Figure S9). In sum, these results support that the subject identification was mainly driven by the neural-origin individual differences rather than head motion.

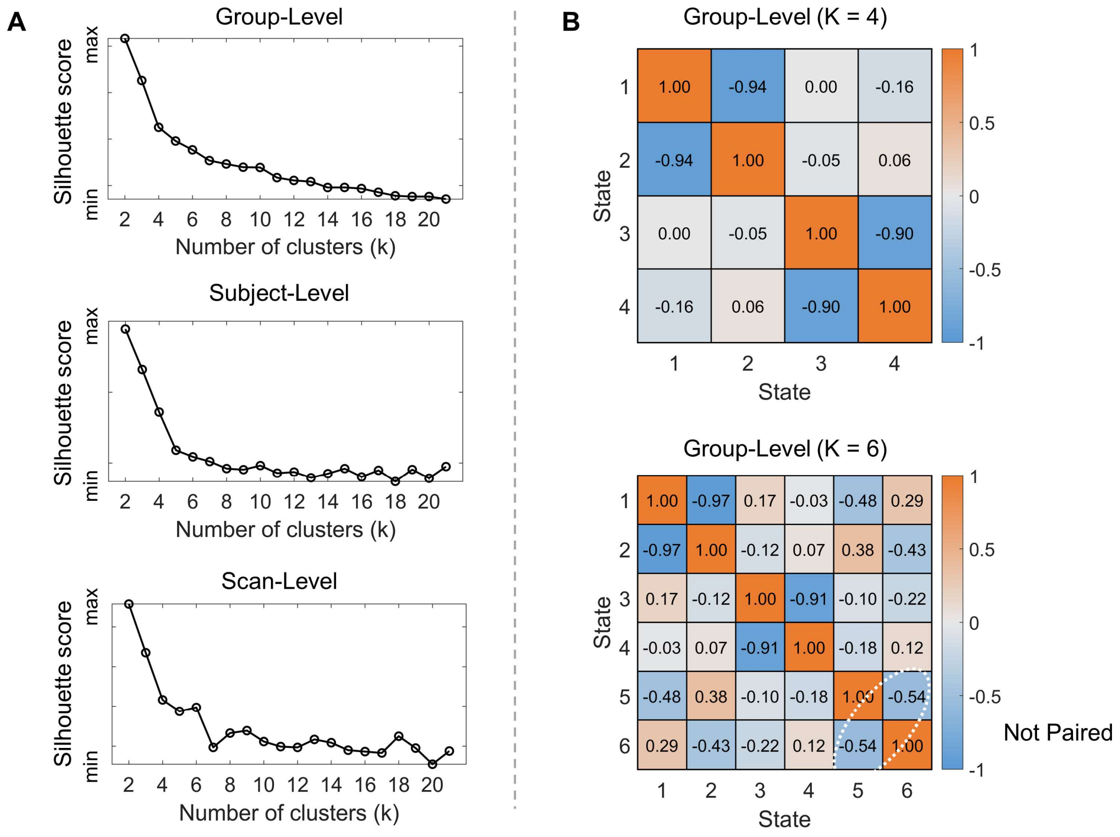


Figure S1. (A) The clustering curve (silhouette score) at the three levels (group-level, subject-level and scan-level). The K-means clustering was performed from K = 2 to K = 21 with step length = 1. Based on the elbow method, we could choose K = 4 or K = 6. (B) The between-state spatial similarity matrix at the group-level. It can be observed that for K = 4, there were two opposite CAP pairs. While for K = 6, there were only two CAP pairs rather than three. Therefore, we analyzed the results of K = 4 in our manuscript.

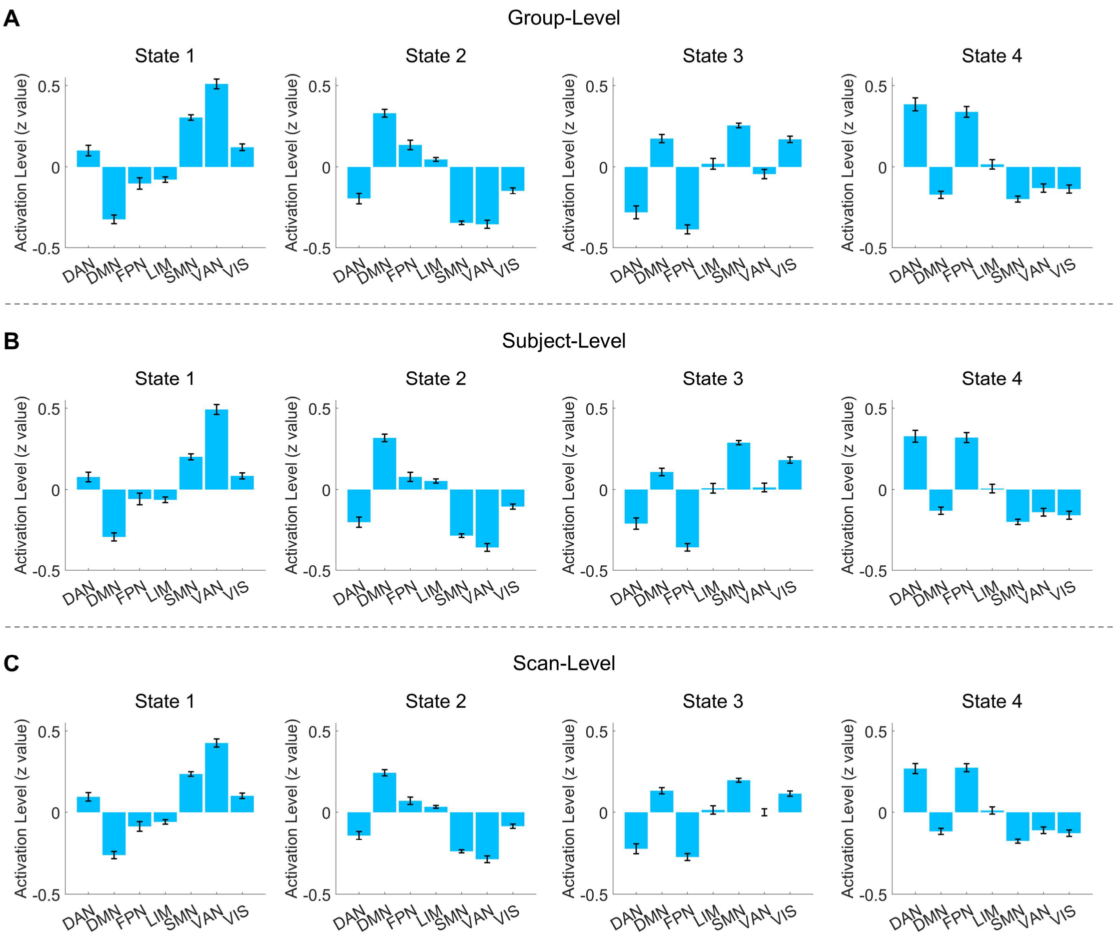


Figure S2. The coactivation level of seven functional networks at the (A) group-level, (B) subject-level and (C) scan-level. The timeseries of each ROI were normalized using Z-score independently. For instance, the activation level of VAN at State 1 was positive, which means the averaged BOLD amplitude of VAN within the time period of State 1 is higher than its baseline over the whole timeseries.

Abbreviations: DAN, dorsal attention network; DMN, default mode network; FPN, fronto-parietal network; LIM, limbic network; SMN, somatomotor network; VAN, ventral attention network; VIS, visual network.

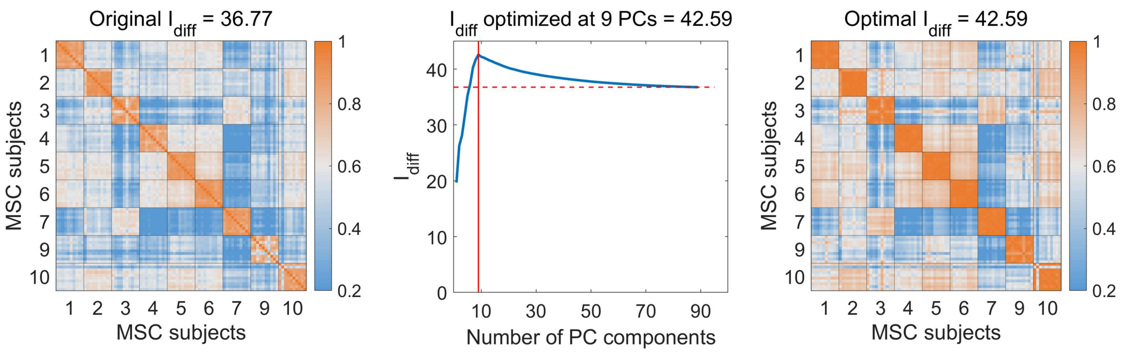


Figure S3. Optimized differential identifiability by using PCA reconstruction at the subject-level, State 3. (A) The original identifiability matrix, $\boldsymbol{I_{diff}}$ = 36.77. (B) Optimal $\boldsymbol{I_{diff}}$ measured using 9 PCs, the red horizontal dashed line indicates the original $\boldsymbol{I_{diff}}$ value. (C) Reconstructed identifiability matrix with maximal $\boldsymbol{I_{diff}}$ = 42.59 using 9 PCs.

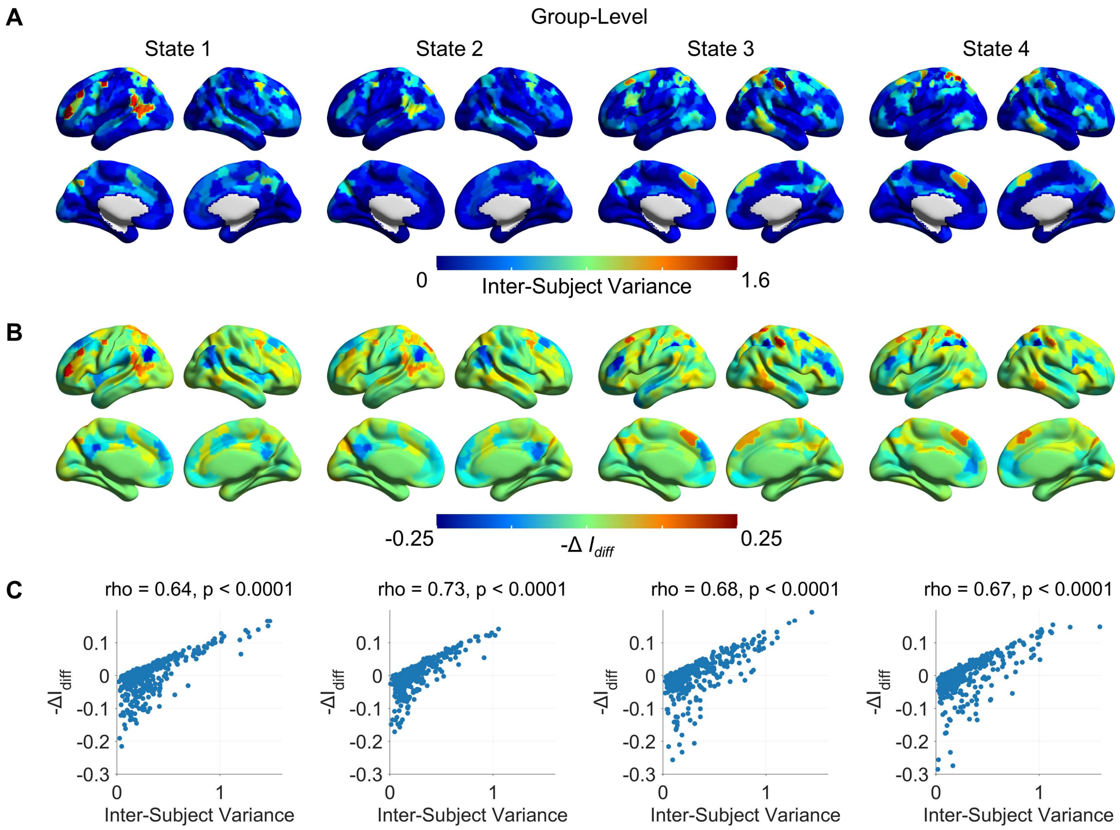


Figure S4. Brain regions contributed to the subject identification at the group-level. (A) Variance maps of the four CAP states. (B) Changed differential identifiability maps. (C) A significant positive correlation was found between the variance map and the changed differential identifiability map.

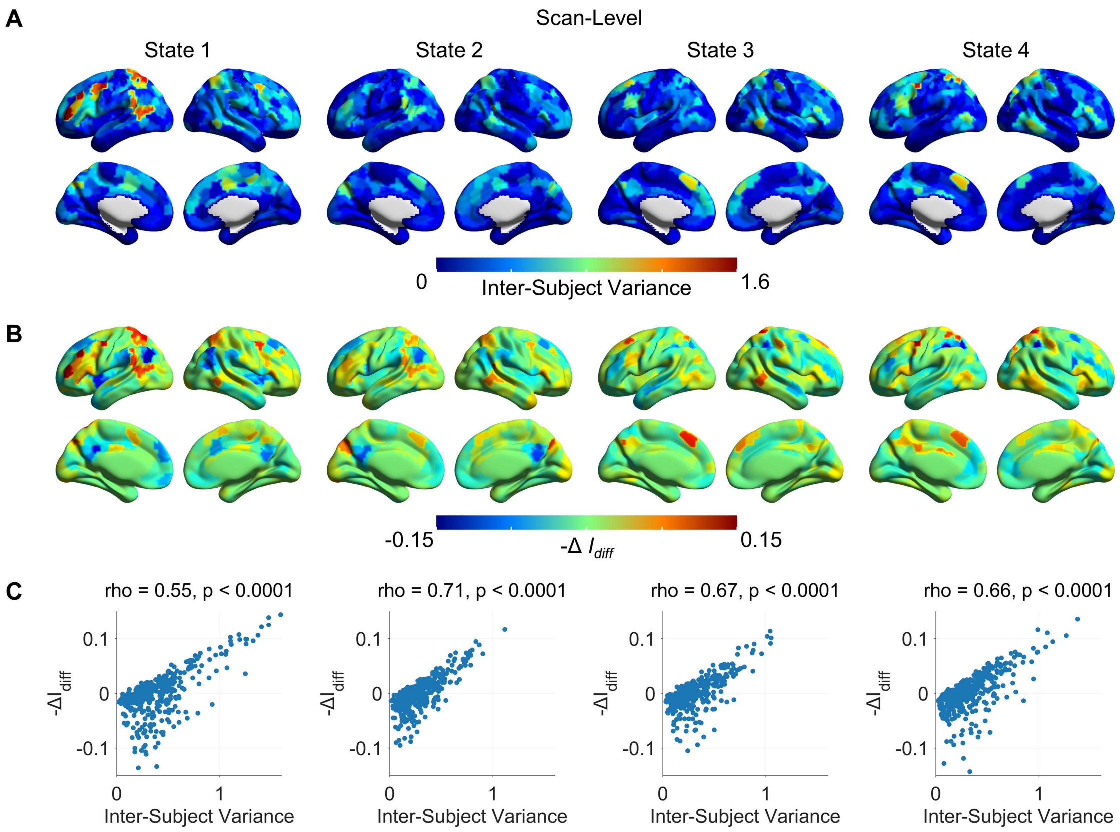


Figure S5. Brain regions contributed to the subject identification at the scan-level. (A) Variance maps of the four CAP states. (B) Changed differential identifiability maps. (C) A significant positive correlation was found between the variance map and the changed differential identifiability map.

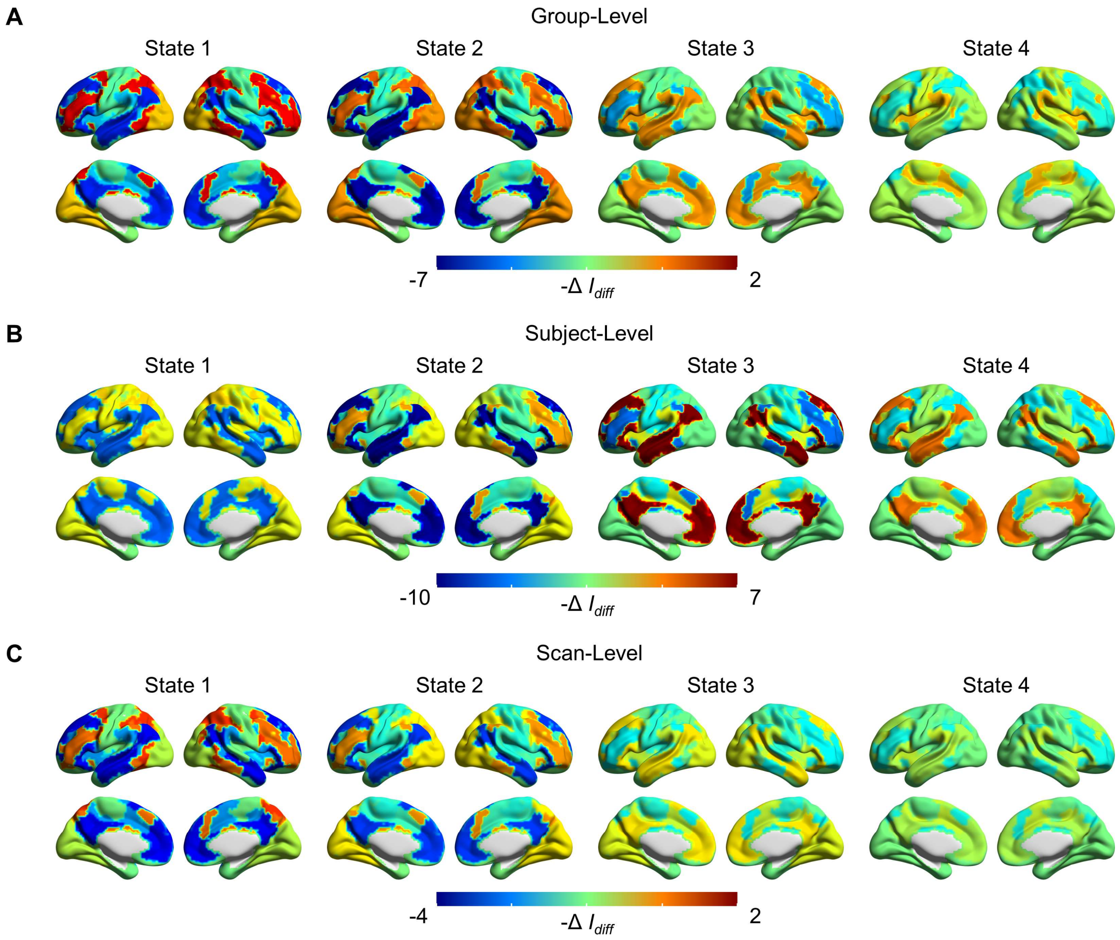


Figure S6. Functional networks contributed to the subject identification at the (A) group-level, (B) subject-level and (C) scan-level. The colorbar indicates the changed differential identifiability. A higher -$\boldsymbol{\Delta I_{diff}}$ value represents a larger contribution.

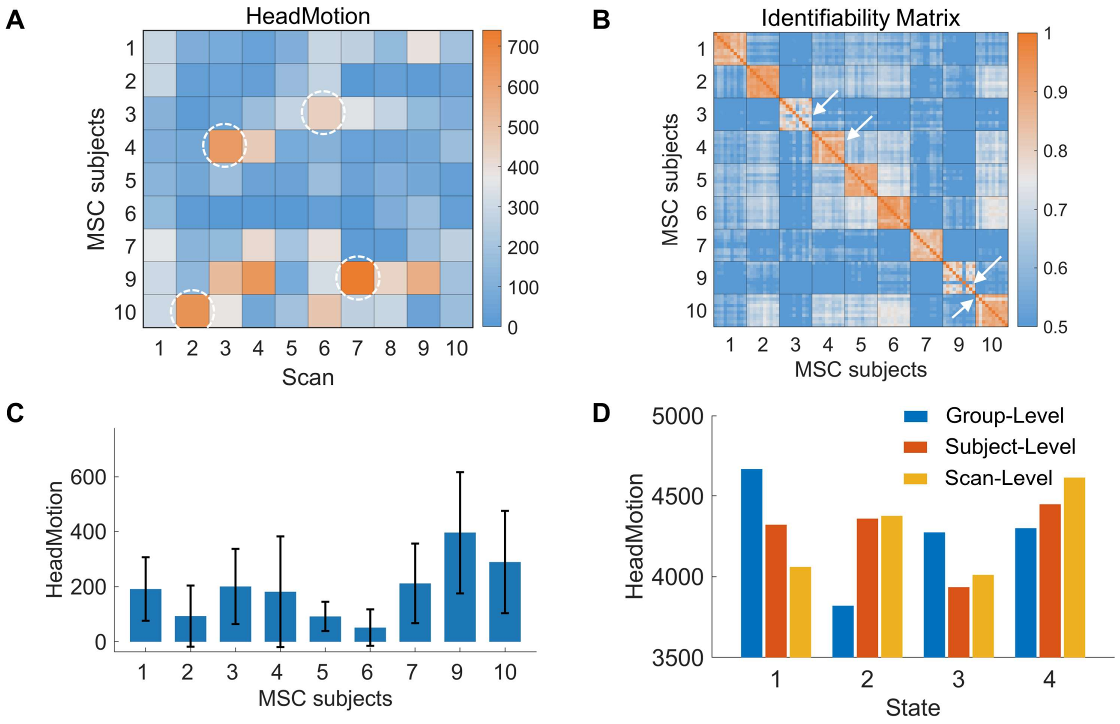


Figure S7. The head motion across scans, subjects and CAP states. (A) Head motion level (number of motion-contaminated frames) across the ninety scans. (B) The identifiability matrix for State 1 at the subject-level. As the white circle and arrow showed, within each individual, the scan with a relatively larger head motion level also showed less spatial similarity with the other scans from the same subject. Motion-contaminated frames of (C) each subject and (D) CAP state. Nine subjects were included in this study, each subject was scanned ten times, and each scan had 818 volumes, hence there were a total of 73,620 (818*9*10) volumes. 17,063 volumes were labeled as motion-contaminated frames if their FD > 0.2 mm. The error bar shows the standard deviation across the ten scans.

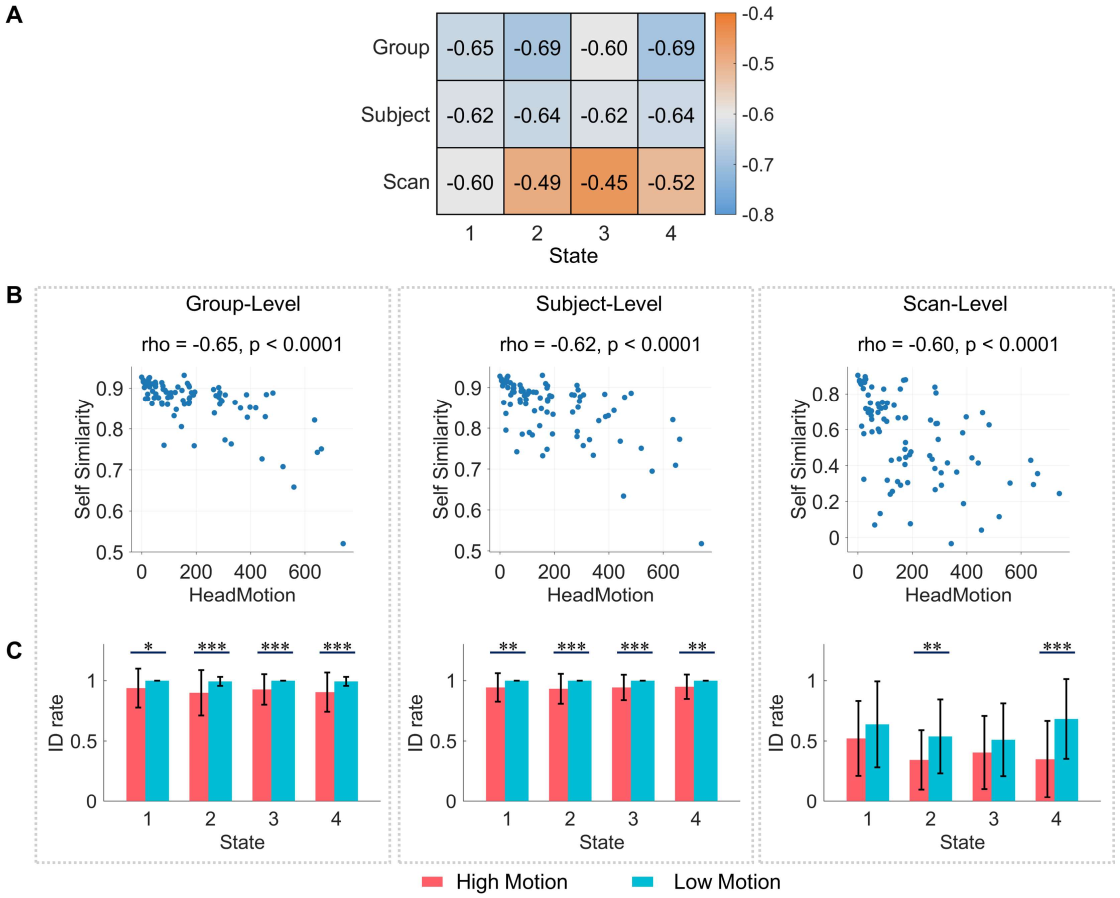


Figure S8. Head motion effects on CAP states and subject identification. (A) The relationship between head motion level and intra-subject similarity. (B) Scatter plots of State 1, each point is a scan, the X-axis is the number of motion-contaminated frames, and the Y-axis shows the self-similarity $I_{self(i,\ j)}$. (C) The identification (ID) rate comparisons between the high motion group and low motion group (top/bottom five scans per subject, ordered by the number of motion-contaminated frames). Permutation test was used to test the group differences, * indicates P < 0.05, ** indicates P < 0.005, and *** indicates P < 0.0005. Errorbar shows the standard deviation.

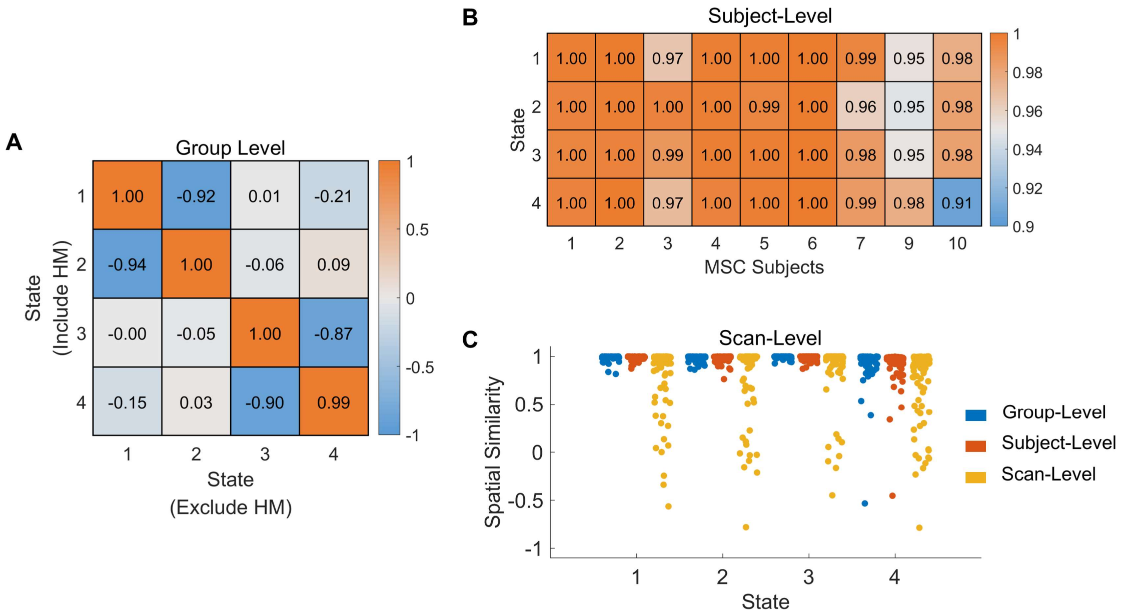


Figure S9. Comparisons between CAP states before and after excluding motion-contaminated frames at the (A) group level, (B) subject-level and (C) scan-level. Particularly, for group-level and subject-level results, their reconstructed CAPs for each scan were also compared in (C).

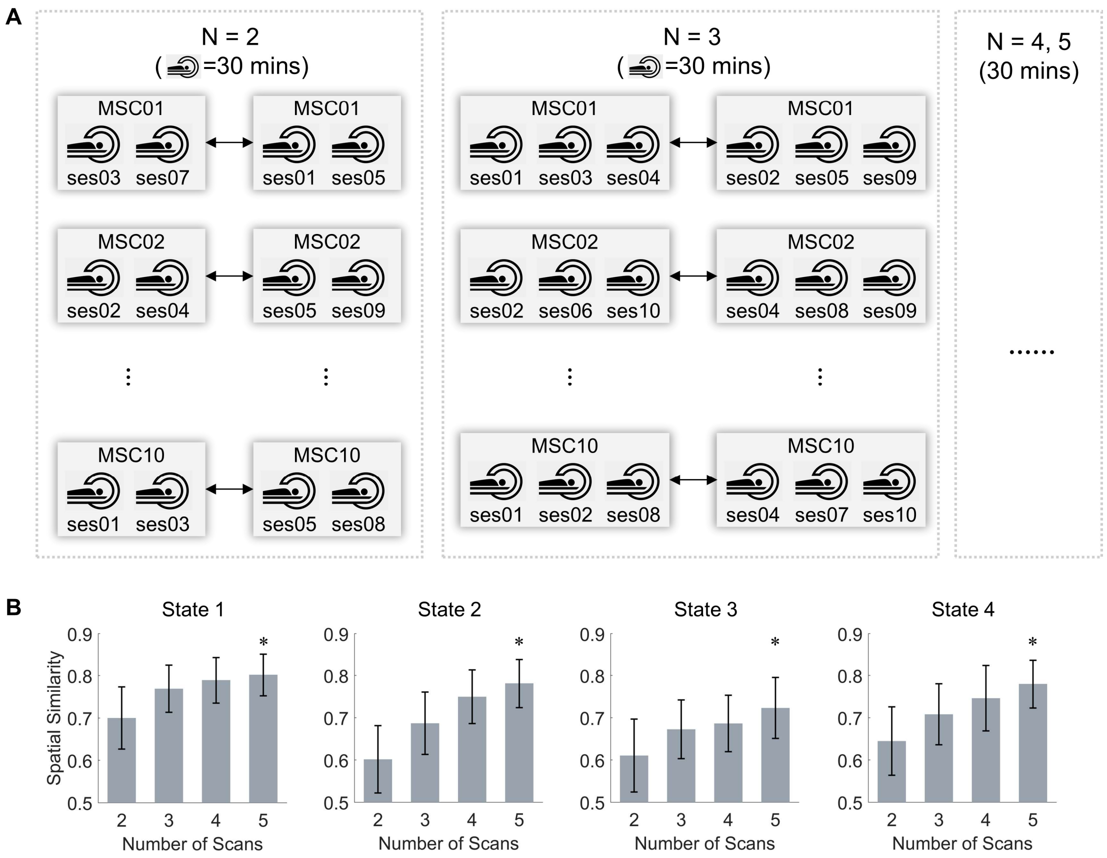


Figure S10. More data per subject increased the generalizability of subject-level CAPs. (A) For each subject, two non-overlapped subgroups with equal scan numbers ($N$ = 2 to 5) were randomly extracted from the ten scans, and the subject-level CAPs were estimated for the two groups independently and compared. (B) The intra-subject spatial similarity for the four CAP states between two subgroups. The intra-subject similarity increased when more scans were included in each subgroup. The spatial similarity was measured by Pearson correlation, and the errorbar shows the standard deviation.

Table S1. The intra-subject similarity, inter-subject similarity and differential identifiability at the three (group-, subject- and scan-) levels.

| **intra-subject similarity ($I_{self}$)** | State 1 | State 2 | State 3 | State 4 |
|---|---|---|---|---|
| group-level | **86.58** | **86.11** | 83.78 | 84.65 |
| subject-level | 85.12 | 85.52 | **83.88** | **84.87** |
| scan-level | 56.66 | 47.06 | 44.43 | 48.15 |
| **inter-subject similarity ($I_{other}$)** | State 1 | State 2 | State 3 | State 4 |
| group-level | **69.60** | **71.04** | **63.93** | **65.61** |
| subject-level | 52.16 | 56.89 | 47.11 | 51.09 |
| scan-level | 40.42 | 34.19 | 30.49 | 33.23 |
| **differential identifiability ($I_{diff}$)** | State 1 | State 2 | State 3 | State 4 |
| group-level | 16.98 | 15.07 | 19.86 | 19.03 |
| subject-level | **32.96** | **28.63** | **36.77** | **33.78** |
| scan-level | 16.23 | 12.88 | 13.94 | 14.93 |

Note: For each CAP state, the largest value among the three levels is bolded.

Table S2. Identification rate at the three (group-, subject- and scan-) levels.

| **Identification Rate (mean±SD)** | State 1 | State 2 | State 3 | State 4 |
|---|---|---|---|---|
| group-level | 0.95 ± 0.11 | 0.94 ± 0.12 | 0.95 ± 0.09 | 0.95 ± 0.09 |
| subject-level | **0.98 ± 0.05** | **0.97 ± 0.06** | **0.97 ± 0.05** | **0.98 ± 0.05** |
| scan-level | 0.57 ± 0.32 | 0.43 ± 0.23 | 0.45 ± 0.26 | 0.52 ± 0.28 |

Note: For each CAP state, the largest value across the three levels is bolded.

Table S3. The optimized differential identifiability by using PCA.

| $I_{diff}$ **(original/optimized)** | State 1 | State 2 | State 3 | State 4 |
|---|---|---|---|---|
| group-level | 16.98 / **19.57** | 15.07 / **17.59** | 19.86 / **23.37** | 19.03 / **22.26** |
| subject-level | 32.96 / **38.52** | 28.63 / **33.18** | 36.77 / **42.59** | 33.78 / **38.97** |
| scan-level | 16.23 / **18.73** | 12.88 / **15.61** | 13.94 / **16.90** | 14.93 / **17.81** |

Note: The first value shows the original $I_{diff}$, and the second value shows the optimized $I_{diff}$.

Table S4. The differential identifiability with/without motion-contaminated frames.

| $I_{diff}$ **(with/without)** | State 1 | State 2 | State 3 | State 4 |
|---|---|---|---|---|
| group-level | 16.98 / **17.16** | **15.07** / 15.04 | **19.86** / 19.48 | **19.03** / 18.62 |
| subject-level | 32.96 / **34.27** | 28.63 / **29.24** | 36.77 / **37.60** | **33.78** / 33.02 |
| scan-level | 16.23 / **18.87** | 12.88 / **13.08** | 13.94 / **15.00** | 14.93 / **16.09** |

Note: The first value shows the $I_{diff}$ with motion-contaminated frames, and the second value shows the $I_{diff}$ without motion-contaminated frames. Larger values are bolded.
